# FGF2 binds to the allosteric site (site 2) and activates integrin αIIbβ3 and FGF1 binds to site 2 but suppresses integrin activation by FGF2: A potential mechanism of anti-thrombotic action of FGF1

**DOI:** 10.1101/2024.04.17.589979

**Authors:** Yoko K Takada, Xueson Wu, David Wei, Samuel Hwang, Yoshikazu Takada

## Abstract

It has been believed that platelet integrin αIIbβ3 recognizes fibrinogen and several ECM proteins, and we recently showed that αIIbβ3 binds to several inflammatory cytokines (e.g., CCL5, and CXCL12), which are stored in platelet granules. These ligands bind to the classical ligand (RGD)-binding site (site 1) of integrin αIIbβ3. Also, they bind to the allosteric site (site 2) of αIIbβ3, which is distinct from site 1, and allosterically activate αIIbβ3. Site 2 is known to be involved in allosteric integrin activation and inflammatory signaling. FGF2 is also stored in platelet granules and known to be pro-thrombotic, but it is unclear if FGF2 binds to αIIbβ3. We studied if FGF2 and its homologue FGF1 bind to αIIbβ3 and induce allosteric activation. FGF1 (not stored in platelet granules) is known to be anti-thrombotic. Mechanism of FGF1’s anti-thrombotic action is unknown. Here we describe that FGF1 and FGF2 bound to site 1 of αIIbβ3, indicating that αIIbβ3 is a new receptor for FGF1/2. Notably, FGF2 bound to site 2 and allosterically activated αIIbβ3.

Point mutations in the site 2-binding interface of FGF2 suppressed this activation, indicating that FGF2 binding to site 2 is required for activation (FGF2 is an agonist to site 2). In contrast, FGF1 bound to site 2 but did not activate αIIbβ3, and instead suppressed integrin activation induced by FGF2, indicating that FGF1 acts as an antagonist of site 2. A non-mitogenic FGF1 mutant (R50E), which is defective in binding to site 1 of αvβ3, suppressed αIIbβ3 activation by FGF2 as effectively as WT FGF1. We propose that FGF1 R50E has therapeutic potential for anti-thrombosis.

## Introduction

Integrins are a superfamily of αβ heterodimers and are receptors for extracellular matrix (ECM) (e.g., vitronectin, fibronectin, and collagen), cell surface proteins (e.g., VCAM-1, and ICAM-1), and soluble ligands (e.g., growth factors) (Takada et al. 2007). Antagonists to integrins have been shown to suppress FGF2-induced angiogenesis and tumor growth, indicating that integrins are required for FGF signaling (FGF-integrin crosstalk) (Brooks et al. 1994). We identified FGF1 and FGF2 as new ligands for integrin αvβ3 by virtual screening of protein data bank (PDB) using docking simulation with the integrin headpiece as a target (Mori et al. 2008, Mori et al. 2017). Docking simulation predicts that FGF1 and FGF2 bind to the classical ligand-binding site of αvβ3 (site 1) (Mori et al. 2008) (Mori et al. 2017). FGF1 binding to integrin αvβ3 induced integrin αvβ3-FGF1-FGFR1 ternary complex formation (Yamaji et al. 2010). The FGF1 mutant defective in integrin binding due to a point mutation in the predicted integrin-binding site (Arg50 to Glu, R50E) did not bind to integrin αvβ3 but still bound to FGFR and heparin and was defective in inducing sustained ERK1/2 activation, ternary complex formation, and inducing mitogenesis (Mori et al. 2008). The R50E FGF1 mutant was defective in mitogenesis and in inducing angiogenesis, and suppressed angiogenesis induced by WT FGF1 (dominant-negative effect) (Mori et al. 2013). We obtained similar result in FGF2 (Mori et al. 2017). The FGF2 mutants (K119E/R120E and K125E) in the integrin-binding interface of FGF2 were defective in signaling, ternary complex formation, and acted as dominant-negative antagonists (Mori et al. 2017). It has been reported that FGF2 and integrin α6β1 are important for maintaining the pluripotency of human pluripotent stem cells (hPSCs). It has recently been reported that integrin α6β1-FGF2-FGFR ternary complex formation is critical for maintaining the pluripotency of hPSCs (Cheng et al. 2024). The FGF2 K125E was incapable of inducing the hPSC properties, such as proliferation, ERK activity, and large focal adhesions at the edges of human induced pluripotent stem cells (hiPSCs) colonies. These findings indicate that FGF1 and FGF2 binding to integrin (site 1) and ternary complex formation is critical for their signaling functions.

αIIbβ3 is known as a receptor for ECM proteins (e.g., fibronectin, fibrinogen, plasminogen, prothrombin, thrombospondin and vitronectin) in addition to CD40L (Andre et al. 2002) (Prasad et al. 2003). We discovered, however, that several proinflammatory proteins such as CX3CL1 (fractalkine) (Fujita et al. 2014), CXCL12 (SDF-1) (Fujita et al. 2018, Takada et al. 2022), Rantes (CCL5) (Takada et al. 2022), and secreted phospholipase A2 type IIA (sPLA2-IIA) (Fujita et al. 2015), CD40L (Takada et al. 2021), and P-selectin (Takada et al. 2023). These findings suggest that αIIbβ3 is as promiscuous as integrin αvβ3 and is affected by inflammatory cytokiens.

Activation of αIIbβ3 is a key event that triggers platelet aggregation upon platelet activation by inducing αIIbβ3 binding to fibrinogen leading to bridge formation between platelets. It has been well established that activation of αIIbβ3 is mediated exclusively by inside-out signaling induced by platelet agonists (e.g., thrombin, ADP, and collagen) (Shattil et al. 2010, Ginsberg 2014). We described that Inflammatory cytokines described above activated αIIbβ3 independent of inside-out signaling. We found that this activation is induced by ligand binding to the allosteric ligand-binding site (site 2) of integrins. Site 2 is distinct from the classical ligand-binding site (site 1) and is on the opposite side of site 1 in the integrin headpiece (Fujita et al. 2014, Fujita et al. 2018, Takada et al. 2022). We showed that peptides from site 2 (site 2 peptides) bound to these allosteric activators and suppressed integrin activation, indicating that they are required to bind to site 2 for allosteric integrin activation. Since the allosteric activation is induced by inflammatory cytokines, there should be a link between allosteric integrin activation and platelet aggregation. Recently, pro-inflammatory lipid mediator, 25-hydroxycholesterol, was found to bind to site 2 of integrins and activates integrins and induce inflammatory signals (e.g., secretion of IL-6 and TNF) (Pokharel et al. 2019), which verify the role of site 2 in allosteric integrin activation and inflammatory signals in inflammation.

Several cytokines, including those stored in platelet granules (CCL5, CXCL12, CD40L, and P-selectin) and rapidly transported to the surface upon platelet activation. These cytokines activate αIIbβ3 by binding to site 2 in an allosteric manner (Takada et al. 2023, Takada et al. 2023). We hypothesize that inflammatory cytokines stored in platelet granules play an important role in activating αIIbβ3. FGF2 is pro-inflammatory and induces the expression of a wide repertoire of inflammation-related genes in endothelial cells, including pro-inflammatory cytokines/chemokines and their receptors, endothelial cell adhesion molecules, and components of the prostaglandin pathway (Presta et al. 2009). FGF2 expression is enhanced in endothelial precursor cells in deep vein thrombosis (Sun et al. 2020), suggesting that FGF2 is pro-thrombotic. FGF2 is stored in platelet granules and rapidly transported to the surface upon platelet activation. It is unclear if FGF2 induces integrin activation by binding to site 2.

FGF1 is another member of the same subfamily as FGF2 (FGF1 subfamily) but is not stored in platelet granules. Previous studies showed that FGF1 is antiinflammatory (Fan et al. 2019); however, the mechanism of anti-inflammatory effects of FGF1 is unclear. Also, FGF1 is shown to lower blood glucose levels in diabetic mice but the mechanism of this action is unknown (Suh et al. 2014). It has been reported that blocking FGF1 synthesis by activating FGF1 promoter methylation exacerbate deep vein thrombosis, suggesting that FGF1 is cardioprotective (anti-thrombotic) (Zhang and Qin 2023). However, the mechanism of the cardioprotective action of FGF1 is unknown.

We hypothesized that FGF2’s pro-thrombotic action and FGF1’s anti-thrombotic action may be mediated by binding to site of αIIbβ3.

We describe here that FGF2 activated αIIbβ3 by binding to site 2. FGF1 also bound to site 2 but did not activate αIIbβ3, and instead suppressed integrin activation induced by FGF2, indicating that FGF1 act as an antagonist of site 2. We propose a model, in which pro-thrombotic action of FGF2 is mediated by αIIbβ3 activation by binding to site 2, and anti-thrombotic action of FGF1 is mediated by inhibiting αIIbβ3 activation by FGF2 by blocking site 2. Non-mitogenic FGF1 R50E also suppressed activation of αIIbβ3 by FGF2 in cell-free conditions. We propose that site 2 of αIIbβ3 may be a novel therapeutic target for thrombotic diseases.

## Materials and Methods

### Materials

Fibrinogen γ-chain C-terminal residues 390-411 cDNA encoding (6 His tagged)[HHHHHH]NRLTIGEGQQHHLGGAKQAGDV] was conjugated with the C-terminus of GST (designated γC390-411) in pGEXT2 vector (BamHI/EcoRI site). The protein was synthesized in E. coli BL21 and purified using glutathione affinity chromatography. The protein was synthesized in E. coli BL21 and purified using glutathione affinity chromatography.

FGF1 (Mori et al. 2008) and FGF2 (Mori et al. 2017) were synthesized as previously described.

Cyclic β3 site 2 peptide fused to GST-The 29-mer cyclic β3 site 2 peptide C260-RLAGIV[QPNDGSHVGSDNHYSASTTM]C288 (C273 is changed to S) was synthesized by inserting oligonucleotides encoding this sequence into the BamHI-EcoRI site of pGEX-2T vector. The positions of Cys residues for disulfide linkage were selected by using Disulfide by Design-2 (DbD2) software (http://cptweb.cpt.wayne.edu/DbD2/) (Craig and Dombkowski 2013). It predicted that mutating Gly260 and Asp288 to Cys disulfide-linked cyclic site 2 peptide of β3 does not affect the conformation of the original site 2 peptide sequence QPNDGSHVGSDNHYSASTTM in the 3D structure. We found that the cyclic site 2 peptide bound to CX3CL1 and sPLA2-IIA to a similar extent to non-cyclized β3 site 2 peptides in ELISA-type assays (data not shown). We synthesized the proteins in BL21 cells and purified using glutathione-Sepharose affinity chromatography.

### Activation of soluble αIIbβ3 by FGF2

ELISA-type binding assays were performed as described previously (Fujita et al. 2018). Briefly, wells of 96-well Immulon 2 microtiter plates (Dynatech Laboratories, Chantilly, VA) were coated with 100 µl 0.1 M PBS containing γC390-411 for αIIbβ3 for 2 h at 37°C. Remaining protein-binding sites were blocked by incubating with PBS/0.1% BSA for 30 min at room temperature. After washing with PBS, soluble recombinant αIbβ3 (1 µg/ml) in the presence or absence of FGF1 or FGF2 was added to the wells and incubated in Hepes-Tyrodes buffer (10 mM HEPES, 150 mM NaCl, 12 mM NaHCO_3_, 0.4 mM NaH_2_PO_4_, 2.5 mM KCl, 0.1% glucose, 0.1% BSA) with 1 mM CaCl_2_ for 1 h at room temperature. After unbound αIbβ3 was removed by rinsing the wells with binding buffer, bound αIbβ3 was measured using anti-integrin β3 mAb (AV-10) followed by HRP-conjugated goat anti-mouse IgG and peroxidase substrates. For the time-course experiments, WT FGF2 and soluble αIbβ3 was incubated for 1–60 min instead of 1 h.

### Activation of integrin αIIbβ3 on the cell surface by FGF2

CHO cells that express recombinant αIIbβ3 (αIIbβ3-CHO) were cultured to nearly confluent in DMEM/10% FCS. Cells were resuspended with DMEM/0.02% BSA and incubated for 30 min at room temperature to block protein-binding sites. Cells were then incubated with WT FGF2 or mutants for 5 min at room temperature and then incubated with FITC-labeled γC390-411 for 15 min at room temperature. Cells were washed with PBS/0.02% BSA and analyzed by FACSCalibur (Becton Dickinson, Mountain View, CA). For blocking experiments, FGF2 was preincubated with Fc-β3 peptide for 30 min at room temperature.

### Docking simulation

Docking simulation of interaction between FGF2 and integrin αvβ3 (closed headpiece form, PDB code 1JV2) was performed using AutoDock3 as described previously (Ieguchi et al. 2010). We used the headpiece (residues 1–438 of αv and residues 55– 432 of β3) of αvβ3 (closed form, 1JV2.pdb). Cations were not present in integrins during docking simulation, as in the previous studies using αvβ3 (open headpiece form, 1L5G.pdb) (Mori et al. 2008, Saegusa et al. 2008). The ligand is presently compiled to a maximum size of 1024 atoms. Atomic solvation parameters and fractional volumes were assigned to the protein atoms by using the AddSol utility, and grid maps were calculated by using AutoGrid utility in AutoDock 3.05. A grid map with 127 × 127 × 127 points and a grid point spacing of 0.603 Å included the headpiece of αvβ3. Kollman ‘united-atom’ charges were used. AutoDock 3.05 uses a Lamarckian genetic algorithm (LGA) that couples a typical Darwinian genetic algorithm for global searching with the Solis and Wets algorithm for local searching. The LGA parameters were defined as follows: the initial population of random individuals had a size of 50 individuals; each docking was terminated with a maximum number of 1 × 10^6^ energy evaluations or a maximum number of 27 000 generations, whichever came first; mutation and crossover rates were set at 0.02 and 0.80, respectively. An elitism value of 1 was applied, which ensured that the top-ranked individual in the population always survived into the next generation. A maximum of 300 iterations per local search were used. The probability of performing a local search on an individual was 0.06, whereas the maximum number of consecutive successes or failures before doubling or halving the search step size was 4.

### Other methods

Treatment differences were tested using ANOVA and a Tukey multiple comparison test to control the global type I error using Prism 10 (Graphpad Software).

## Results

### FGF1 and FGF2 bind to the classical binding site (site 1) of αIIbβ3

Previous studies showed that pro-inflammatory proteins are stored in platelet granules (e.g., CCL5, CXCL12, CD40L, and P-selectin) and rapidly transported to the surface upon platelet activation. These cytokines bind to site 1 of integrin (See Introduction). We showed that FGF1 and FGF2 bound to site 1 of integrin αvβ3 (Mori et al. 2008) (Mori et al. 2017), leading to the αvβ3-FGF1/2-FGFR ternary complex on the cell surface. FGF2 is stored in platelet granules and rapidly transported to the surface upon platelet activation (FGF1 is not store in platelet granules), but it is unknown if FGF2 or FGF1 binds to αIIbβ3. We studied if soluble αIIbβ3 bound to FGF2 and FGF1 in ELISA-type binding assays. Wells of 96-well microtiter plate were coated with FGF2 or FGF1 and incubated with soluble αIIbβ3 (ectodomain) in 1 mM Mn^2+^ (Fig. 1a and 1b). We detected the binding of soluble αIIbβ3 to FGF1 and FGF2 in a dose-dependent manner. To show the specificity of FGF2 and FGF1 binding to αIIbβ3, we tested if FGF2 and FGF1 compete with known ligand for αIIbβ3 for binding to αIIbβ3. The disintegrin domain of ADAM-15, a specific ligand for αIIbβ3 (Langer et al. 2005), potently suppressed FGF2 and FGF1 binding to αIIbβ3 (Fig. 1c and 1d), indicating that the binding of FGF2 and FGF1 to αIIbβ3 is specific. FGF2 is known to bind to site 1 of αvβ3 (Mori et al. 2017), but it is unclear if FGF2 binds to αvβ3 and αIIbβ3 in a similar manner. We previously reported that FGF2 mutants defective in binding to site 1 of αvβ3 (the K119E/R120E and K125E mutants) (Mori et al. 2017). The FGF2 mutants defective in binding to αvβ3 bound to αIIbβ3 in 1 mM Mn^2+^ (Fig 1e), indicating that FGF2 binds to site 1 of αIIbβ3 and αvβ3 in a different manner.

**Fig. 1.**
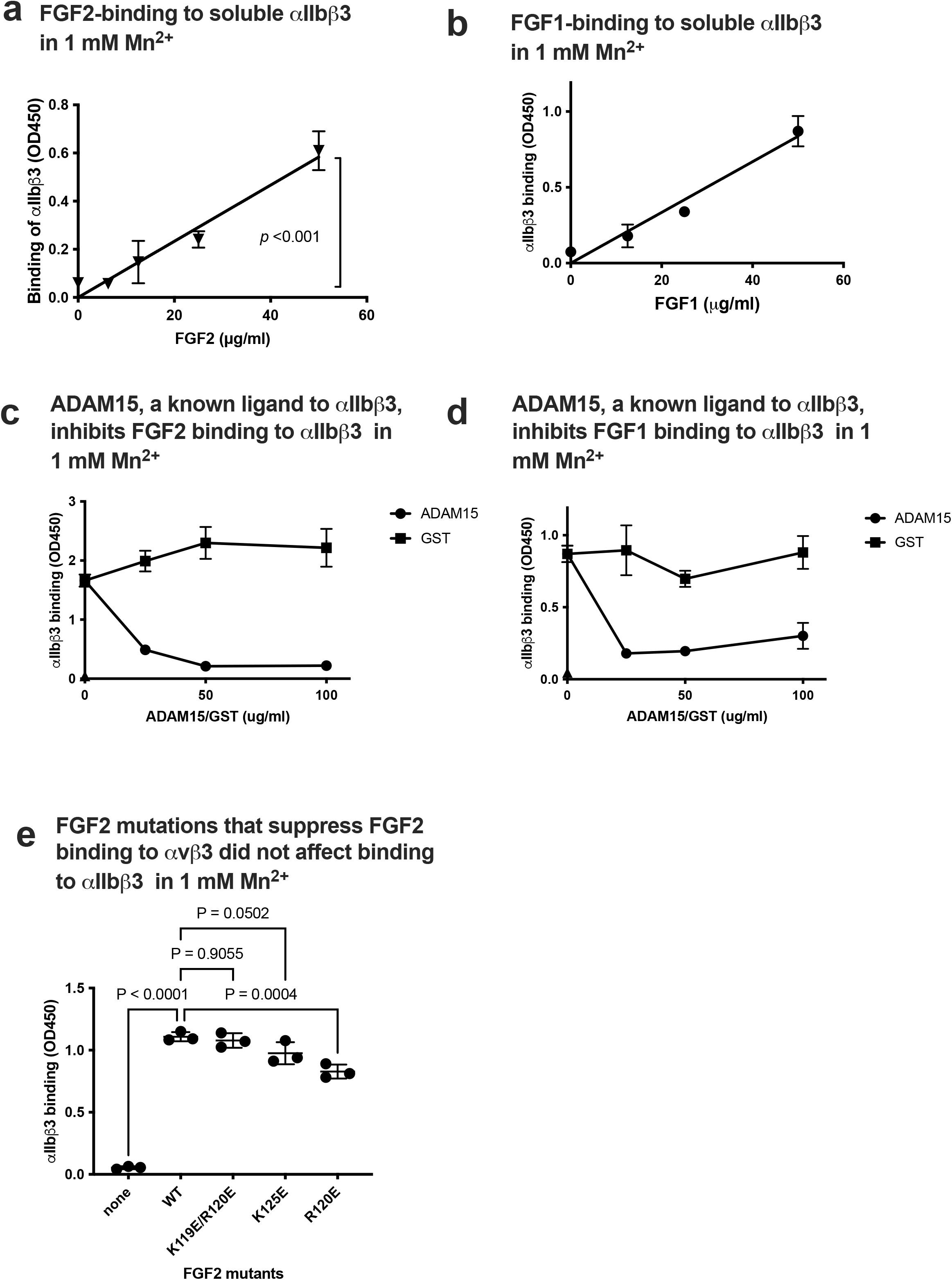
FGF2 and FGF1 bind to site 1 of αIIbβ3. (a) FGF2 binds to soluble αIIbβ3. Wells of 96-well microtiter plate were coated with FGF2 and incubated with soluble αIIbβ3 in 1 mM Mn^2+^. Bound αIIbβ3 was quantified using anti-β3 and HRP-conjugated anti mouse IgG. The data is shown as means +/-SD in triplicate experiments. (b) FGF1 binds to soluble αIIbβ3. Binding of soluble αIIbβ3 to FGF1 was measured as in (a) except FGF1 was used instead of FGF2. The data is shown as means +/-SD in triplicate experiments. (c and d) Inhibition of FGF1/FGF2 binding to soluble αIIbβ3 by ADAM15. Wells of 96-well microtiter plate were coated with FGF2 (c) or FGF1 (d) and incubated with soluble αIIbβ3 in the presence of ADAM-15 disintegrin domain fused to GST or control GST in 1 mM Mn^2+^. The data is shown as means +/-SD in triplicate experiments. (e) Effect of FGF2 mutation that blocked binding to αvβ3 (site 1) on binding to αIIbβ3. FGF2 mutants defective in binding to αvβ3 (site 1) were tested their ability to bind to αIIbβ3 in 1 mM Mn^2+^. The data is shown as means +/-SD in triplicate experiments.

### FGF2 activates soluble integrin αIIbβ3 by binding to the allosteric site (site 2)

It has been established that αIIbβ3 is activated exclusively by inside-out signaling induced by platelet agonists (e.g., thrombin and ADP) (Ginsberg 2014) (Shattil et al. 2010). Previous studies showed, however, that CCL5, CXCL12, CD40L, and CD62P platelet granular contents, activated αIIbβ3 in an allosteric manner by binding to site 2, independent of inside-out signaling (Takada et al. 2023) (Takada et al. 2023) (Takada et al. 2022). We hypothesized that inflammatory cytokines stored in platelet granules activate αIIbβ3 upon platelet activation. It is unknown if FGF2 binds to site 2 and activates αIIbβ3. We studied if αIIbβ3 can be activated by FGF2.

We performed docking simulation of interaction between closed headpiece integrin αvβ3 (1JV2.pdb) and FGF2 using Autodock3. The 3D structure of closed headpiece αvβ3 structure (1JV2.pdb) was used instead of αIIbβ3, since closed headpiece conformation is well defined in αvβ3, but not in αIIbβ3. The simulation predicts that FGF2 binds to site 2 of αvβ3 (docking energy −20.5 kcal/mol) (Fig. 2a). [docking simulation with αIIbβ3, −22.95 kcal/mol]

**Fig. 2.**
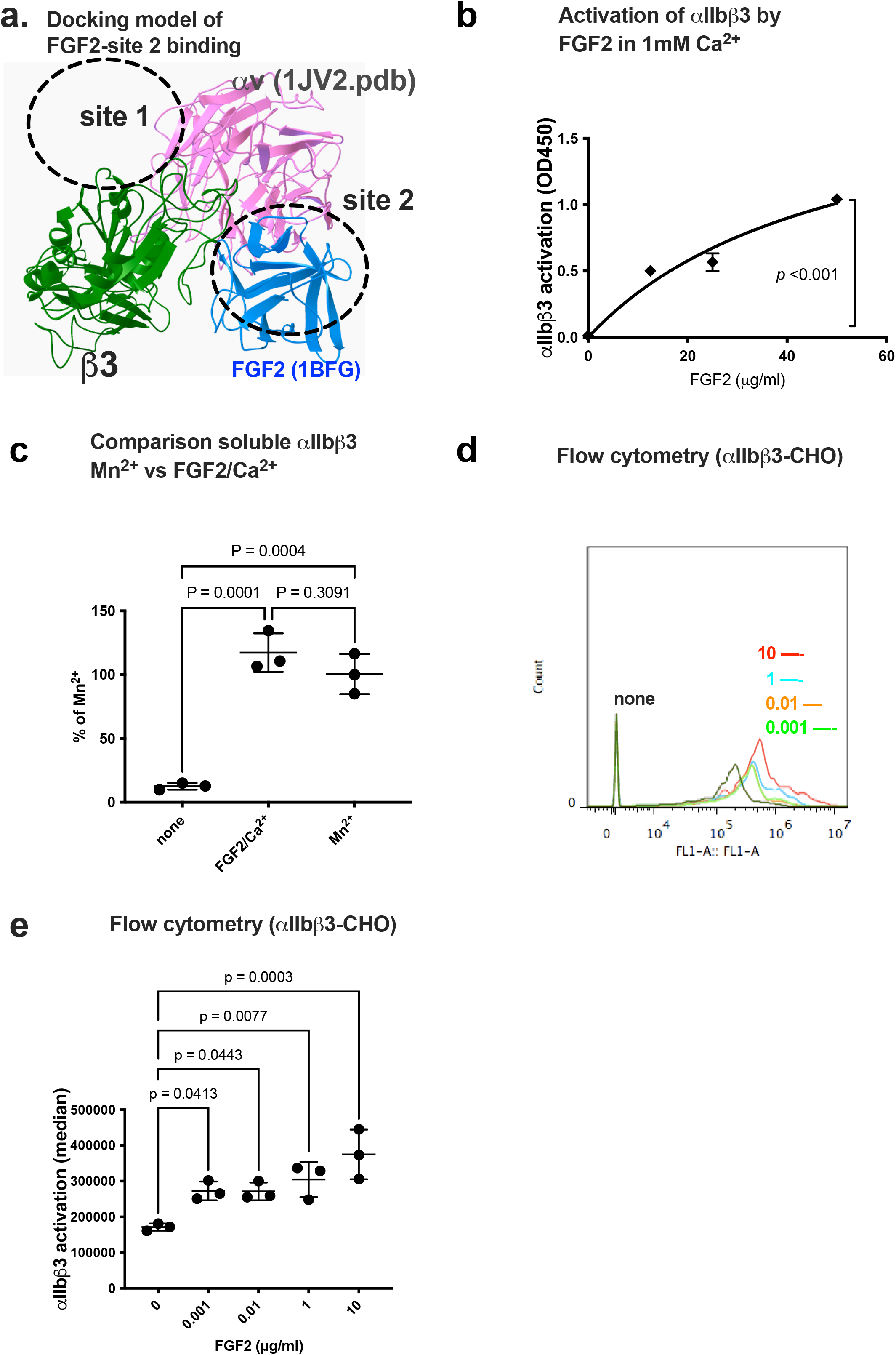
FGF2 binds to site 2 of integrin αIIbβ3 and activates αIIbβ3 (act as an agonist). (a) Docking simulation of interaction between FGF2 (1BFG.pdb) and αvβ3 (closed headpiece form, 1JV2.pdb). FGF2 was also predicted to bind to site 2. (b) FGF2 activated αIIbβ3. The fibrinogen fragment (γC390-411), a specific ligand to αIIbβ3, was immobilized to wells of 96-well microtiter plate and incubated with soluble αIIbβ3 (1 µg /ml) and bound αIIbβ3 was measured in 1 mM Ca^2+^ (to keep integrins inactive). The data is shown as means +/-SD in triplicate experiments. (c). FGF2 and 1 mM Mn^2+^ activate soluble αIIbβ3 to a similar extent. Activation of αIIbβ3 was measured as described in (b) using 1 mM Mn^2+^ or FGF2 (50 µg/ml). (d) and (e). FGF2 activates cell-surface αIIbβ3. CHO cells that express recombinant αIIbβ3 (αIIbβ3-CHO cells) were incubated with FITC-labelled γC390-411, a specific ligand for αIIbβ3, in the presence of FGF2 and bound γC390-411 was measured in flow cytometry.

We found that FGF2 activated integrin αIIbβ3 in 1 mM Ca^2+^ in cell-free conditions in ELISA-type integrin activation assays. Wells of 96-well microtiter plate were coated with a fibrinogen fragment (γC390-411), a specific ligand for αIIbβ3, and incubated with soluble integrin αIIbβ3 in 1 mM Ca^2+^ in the presence of FGF2. We found that FGF2 enhanced the binding capacity of soluble αIIbβ3 in a dose-dependent manner (Fig. 2b).

It has been assumed that 1 mM Mn^2+^ fully activates integrins (Gailit and Ruoslahti 1988) (Mould et al. 1995) (Altieri 1991) (Elices et al. 1991). We compared the levels of activation of soluble αIIbβ3 by FGF2 with that of 1 mM Mn^2+^ as a standard integrin activator. The level of activation of αIIbβ3 by FGF2 was comparable to that of 1 mM Mn^2+^ (Fig. 2c).

CHO cells that express recombinant αIIbβ3 (αIIbβ3-CHO) have been widely used as a model of αIIbβ3 activation. αIIbβ3 on CHO cells is not activated by platelet agonists (e.g., ADP and thrombin) since CHO cells do not have machinery for inside-out signaling (Ginsberg 2014) (Shattil et al. 2010). We showed that αIIbβ3 on CHO cells was activated by FGF2 (Fig. 2d and 2e), which is consistent with the idea that FGF2 activates αIIbβ3 independent of inside-out signaling. Previous studies showed that several integrin ligands that bind to site 2 and activates integrins bound to peptides from site 2 (of β1 and β3) (Fujita et al. 2014, Fujita et al. 2015, Fujita et al. 2018, Takada et al. 2021, Takada et al. 2022, Yoko K Takada 2022). We found that cyclic peptides derived from site 2 of β3 bound to FGF2 at higher levels than control scrambled β3 peptide, indicating that FGF2 is required to bind to site 2 to activate αIIbβ3 (paper submitted).

### Point mutations of the predicted site 2-binding interface in FGF2 block FGF2-mediated activation of αIIbβ3

To further show how FGF2 binds to site 2 of αIIbβ3, we generated FGF2 mutants defective in site 2 binding. The docking simulation of interaction between FGF2 (1BFG.pdf) and αvβ3 (1JV2.pdb) predicts that amino acid residues Lys66, Arg72, Lys77, Lys86, and Lys110 of β3 interact with αvβ3 (docking energy −20.5 Kcal/mol) (Fig. 3a).

**Fig. 3.**
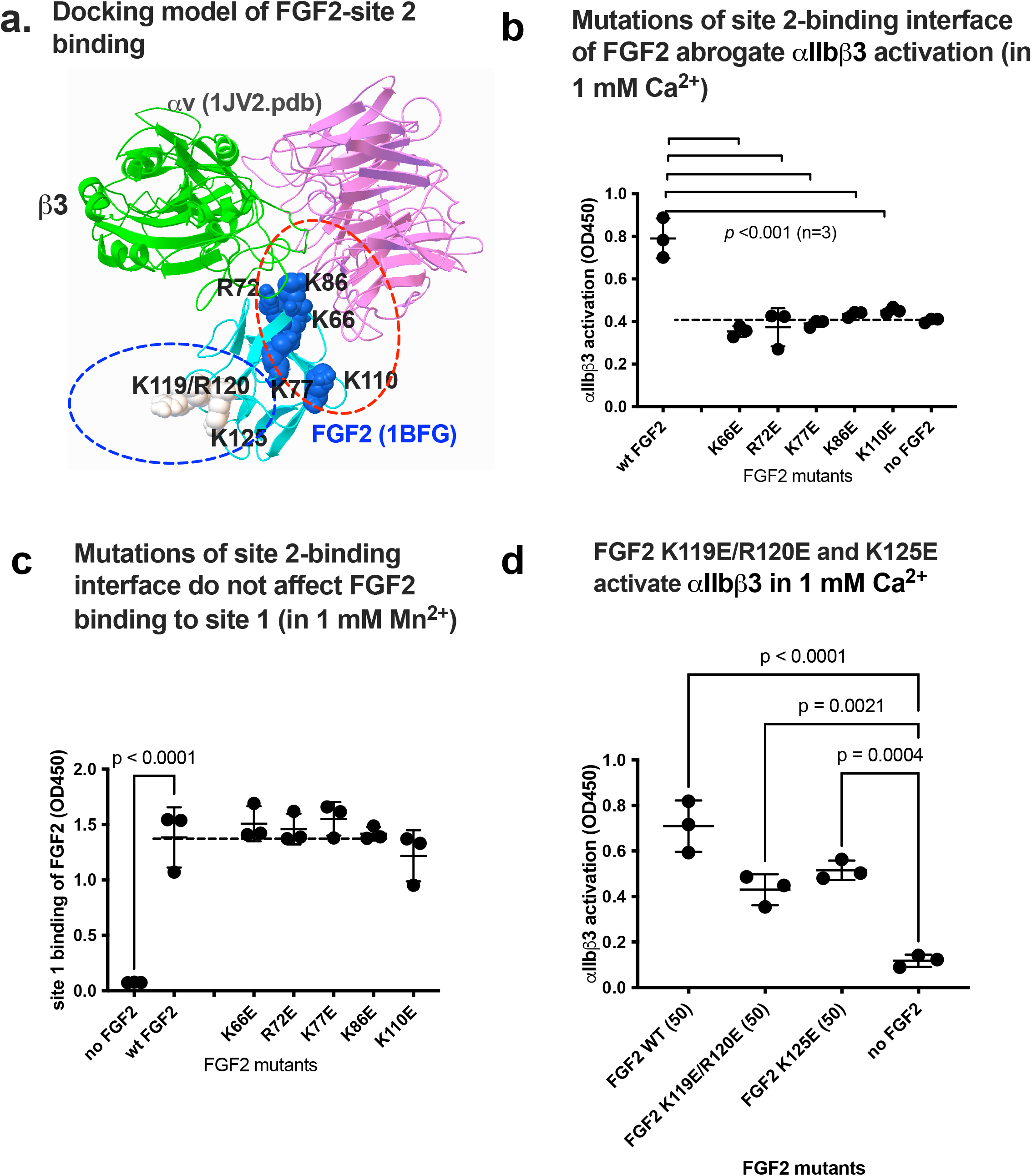
Point mutations in site 2-binding interface of FGF2 effectively reduce activation of integrin αIIbβ3 by FGF2. (a) Positions of amino acid residues involved in site 2 binding predicted by docking simulation. We previously reported that K119E/R120E and K125E mutations suppressed FGF2 binding to integrin site 1 of αvβ3 and thereby suppressed FGF2 mitogenicity (Mori et al. 2017). Arg72, Lys77, Lys86 and Lys110 are in the predicted site 2-binding interface of FGF2. (b) Mutations in the site 2 binding interface of FGF2 blocked activation of αIIbβ3 in 1 mM Ca^2+^. These mutations also blocked activation of αvβ3. The data is shown as means +/-SD in triplicate experiments. c. The point mutations in the predicted site 2-binding site of FGF2 did not affect FGF2 binding to site 1 in 1 mM Mn^2+^. The results indicate that site 1 and site 2-binding sites in FGF2 is distinct and is consistent with the observation that K119E/R120E and K125E can activate integrins. The data is shown as means +/-SD in triplicate experiments. d. Mutations of K119E/R120E and K125E were not effective in blocking allosteric activation by FGF2 by binding to site 2. Note these mutations are not present in the site 2-binding interface (Fig. 6a). The data is shown as means +/-SD in triplicate experiments.

We found that mutation of these amino acids to Glu (K66E, K72E, K77E, K86E, and K110E mutations) suppressed integrin activation by FGF2 (Fig. 3b), indicating that the docking model is correct and that FGF2 binding to site 2 is required for activation of αIIbβ3. We studied if FGF2 mutants defective in binding to site 2 bind to site 1. The wells of 96-well microtiter plate were coated with FGF2 (wild-type or mutants) and incubated with soluble αIIbβ3 in 1 mM Mn^2+^ and bound αIIbβ3 was quantified using anti-β3. We found that K66E, K72E, K77E, K86E, and K110E mutations did not block FGF2 binding to site 1 of αIIbβ3 in 1 mM Mn^2+^ (Fig. 3c), indicating that the effect of the mutations in the site 2-binding site is specific.

We previously showed that FGF2 binds to the classical ligand-binding site (site 1) and the FGF2 mutants (K119E/R120E and K125E) in the integrin-binding interface of FGF2 were defective in signaling, ternary complex formation, and acted as dominant-negative antagonists (Mori et al. 2017). We studied if FGF2 mutants defective in site 1 binding activate αIIbβ3. The K119E/K120E and K125E mutants defective in site 1 binding activated αIIbβ3 (Fig. 3d), indicating that the site 2 and site 1-binding sites in FGF2 are distinct. These findings suggest that allosteric activation of αIIbβ3 requires the binding of FGF2 to site 2 of αIIbβ3.

### FGF1 binds to site 2 but does not activate integrin αIIbβ3 and suppresses αIIbβ3 activation induced by FGF2

Docking simulation of interaction between FGF1 and closed headpiece αvβ3 (1JV2.pdb) predicts that FGF1 binds to site 2 of αvβ3 (docking energy −20.1 kcal/mol) (Fig. 4a).

**Fig. 4.**
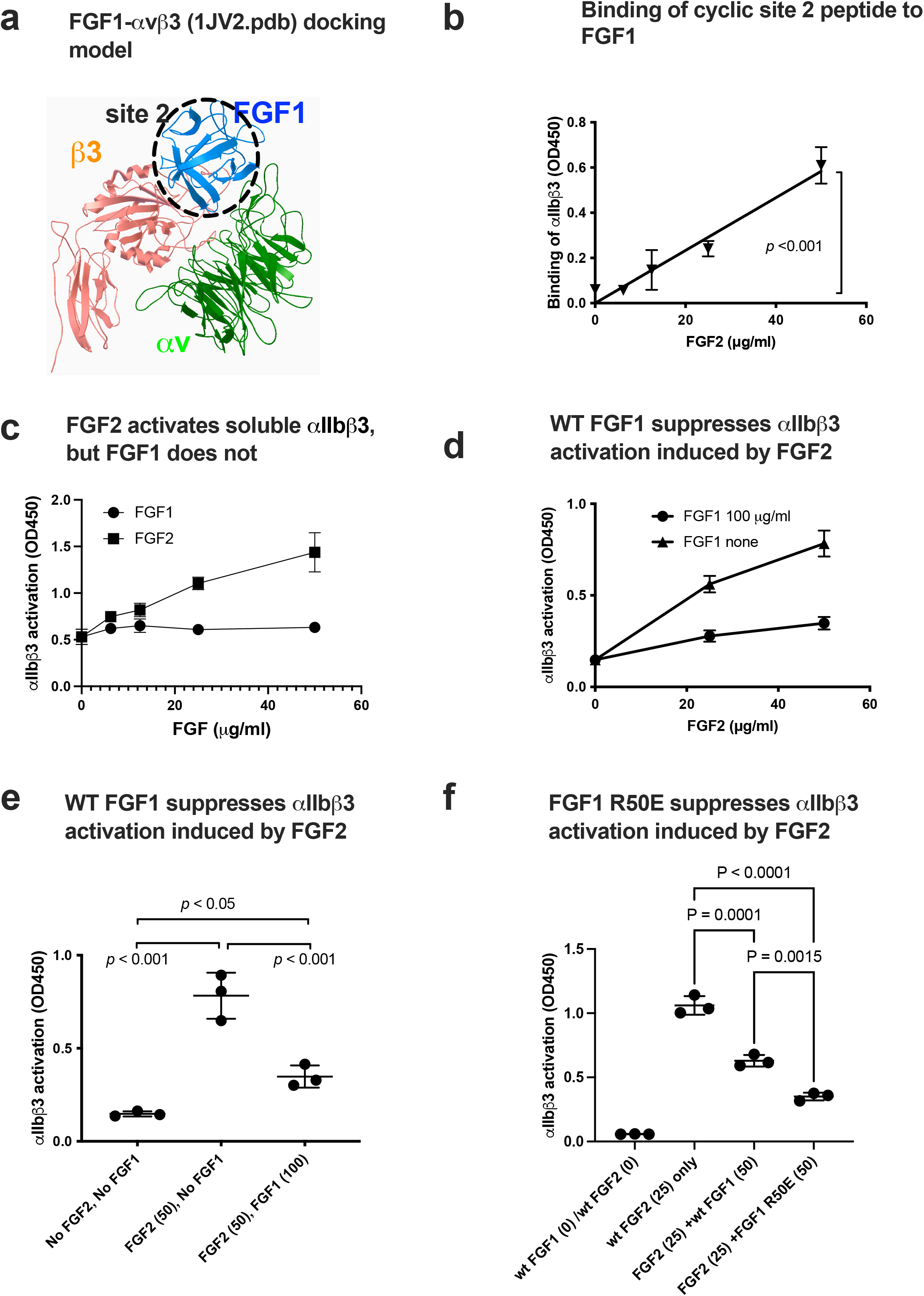
FGF1 is predicted to bind to soluble integrin αIIbβ3 but FGF1 effectively suppresses FGF2-induced activation of integrin. (a) FGF1 is predicted to bind to site 2. Docking simulation of interaction between FGF1 and site 2 of close headpiece form of αvβ3 (1JV2.pdb). (b) FGF1 does not activate soluble αIIbβ3. Wells of 96-well microtiter plate were coated with γC390-411, a specific ligand to αIIbβ3, and incubated with soluble αIIbβ3 in the presence of WT FGF2 or FGF1 in 1 mM Ca^2+^. The data is shown as means +/-SD in triplicate experiments. (c) Binding of cyclic site 2 peptide to FGF1. (d) and (e). FGF1 suppresses FGF2-induced activation of soluble αIIbβ3. Activation of soluble αIIbβ3 was assayed as described in (b). The data is shown as means +/-SD in triplicate experiments.

**Fig. 5.**
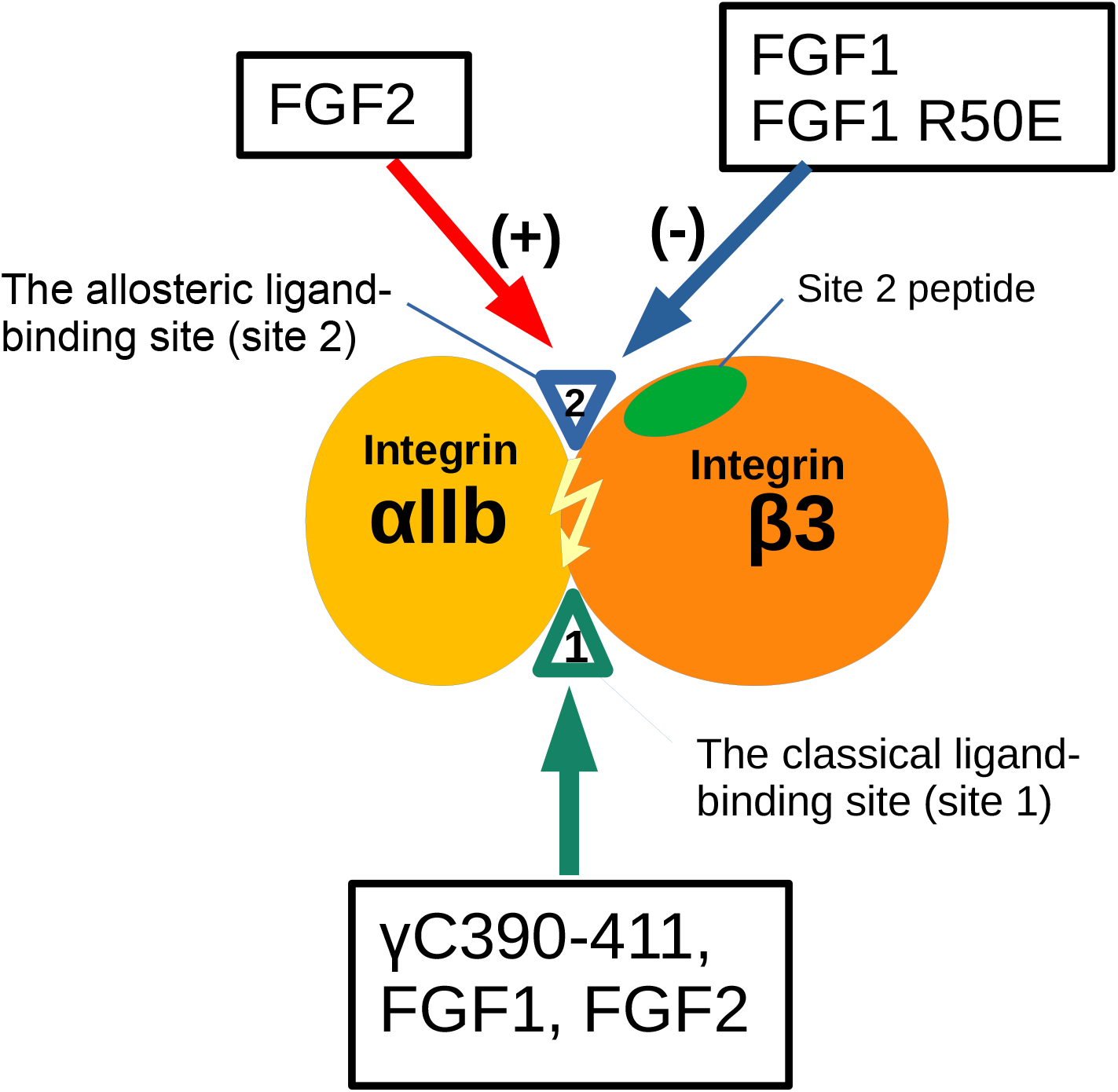
Agonistic action of FGF2 and antagonistic action of FGF1 to the allosteric site (site 2) of β3 integrins. Fibrinogen γ-chain C-terminal peptide (γC390-411) specifically binds to the classical ligand binding site (site1) of αIIbβ3. The present study showed that FGF1 and FGF2 bind to the site1 of αIIbβ3. We showed that FGF2 (stored in platelet granules) binds to site 2 as well and induces allosteric αIIbβ3 activation, leading to platelet aggregation. In contrast, FGF1 binds to site 2 but suppresses integrin activation of αIIbβ3 induced by FGF2. This is a potential mechanism of anti-thrombotic action of FGF1. Non-mitogenic FGF1 R50E mutant is comparable to WT FGF1 in inhibiting FGF2-induced αIIbβ3 activation. FGF1 R50E has potential as an anti-thrombotic agent.

Consistently, cyclic site 2 peptide bound to FGF1 in a dose-dependent manner (Fig. 4b). Thus, we expected that FGF1 also activates αIIbβ3. Unexpectedly, FGF1 did not activate αIIbβ3 at all under the conditions in which FGF2 activated αIIbβ3 (Fig. 4c).

Notably, FGF1 suppressed αIIbβ3 activation by FGF2 (Fig. 4d, 4e), indicating that FGF1 binds to site 2 and acts as an inhibitor of allosteric activation (site 2 antagonist).

We showed that FGF1 R50E is defective in inducing mitogenic signals (e.g., sustained ERK1/2 activation and AKT activation) and acted as an antagonist of FGF1-induced mitogenesis, although R50E still bound to FGFR. FGF1 R50E suppressed angiogenesis and tumor growth in vivo (Mori et al. 2013). We found that FGF1 R50E suppressed FGF2-induced αIIbβ3 activation, as effectively as WT FGF1 (Fig. 4f).

We needed high concentrations of FGF2 and FGF1 for activation or inhibition of activation of soluble integrin in cell-free conditions. Since FGF2 and FGF1 bind to proteoglycans, FGF2 and FGF1 can be highly concentrated on the cell surface, activation or inhibition of activation of integrins is biologically relevant.

## Discussion

It has been believed that platelet integrin αIIbβ3 is more or less specific to fibrinogen and several extracellular matrix (ECM) proteins. We recently showed that αIIbβ3 binds to several pro-inflammatory proteins (e.g., CX3CL1, CXCL12, CCL5 (Takada et al. 2022), CD40L (Takada et al. 2023) and CD62P (Takada et al. 2023)). The present study establishes that FGF1 and FGF2 bind to site 1 of αIIbβ3, indicating that αIIbβ3 is a new receptor for FGF1 and FGF2. These findings suggest that αIIbβ3 is as promiscuous as integrin αvβ3.

Activation of αIIbβ3 is a key event in platelet aggregation and subsequent thrombus formation. We found that FGF2 bound to site 2 and allosterically activated αIIbβ3. We propose that several inflammatory proteins (FGF2, CX3CL1, CXCL12, CCL5, CD40L, and CD62P) in platelet granules play a critical role in allosteric αIIbβ3 activation and subsequently induce platelet aggregation and thrombus formation. The present study suggests that FGF2 may be involved in in platelet aggregation by triggering αIIbβ3 activation in an allosteric manner. These findings are consistent with the recent report that FGF2 expression is enhanced in endothelial precursor cells in deep vein thrombosis (Sun et al. 2020). It is likely that inflammatory cytokines stored in platelet granules, including FGF2, may play a critical role in triggering platelet aggregation activating αIIbβ3 in an allosteric manner.

FGF1 bound to site 2 of αIIbβ3, but did not activate αIIbβ3. Instead, FGF1 suppressed activation of αIIbβ3 induced by FGF2, indicating that FGF1 acts as an antagonist of site 2 (Fig. 8a). FGF1 is shown to be cardioprotective (anti-thrombotic) and blocking FGF1 synthesis by activating FGF1 promoter methylation exacerbate deep vein thrombosis (Zhang and Qin 2023). However, the mechanism of the anti-thrombotic action of FGF1 is unclear. We propose that FGF1’s anti-thrombotic action may be mediated by blocking αIIbβ3 activation by FGF2 from platelet granules. Therefore, we will need to study inflammatory signaling through site 2 in more detail in future studies.

There is growing recognition of the critical role of platelets in inflammation and immune responses (Zaid and Merhi 2022) (Wienkamp et al. 2022). Recent studies have indicated that anti-platelet medications may reduce mortality from infections and sepsis, which suggests possible clinical relevance of modifying platelet responses to inflammation. Platelets release numerous inflammatory mediators that have no known role in hemostasis. Many of these mediators modify leukocyte and endothelial responses to a range of different inflammatory stimuli. Additionally, platelets form aggregates with leukocytes and form bridges between leukocytes and endothelium, largely mediated by platelet CD62P. Thus, the present finding that pro-inflammatory FGF2 allosterically activates αIIbβ3 suggests that FGF2 stored in platelet granules also plays a role in inflammatory actions of platelets. Also, FGF1 may exert anti-inflammatory action by suppressing αIIbβ3 activation in platelets and subsequently blocking platelet activation and thrombosis.

The present study showed that FGF2 bound to site 2 and activated αIIbβ3, as predicted, indicating that FGF2 acts as an agonist of site 2 (Fig. 8b). However, FGF1 bound to site 2 of αIIbβ3, but did not activate αIIbβ3. Instead, FGF1 suppressed activation of αIIbβ3 induced by FGF2, indicating that FGF1 acts as an antagonist of site 2. It has been well established that FGF1 is anti-inflammatory, but the mechanism of anti-inflammatory action of FGF1 is unknown. Previous studies showed that FGF1 remarkably lowered levels of several serum inflammatory cytokines and impeded the inflammatory response (Suh et al. 2014, Liang et al. 2018). FGF1 significantly prevented the development of nonalcoholic fatty liver disease (NAFLD) and diabetic nephropathy (DN) (Liu et al. 2016, Liang et al. 2018). These findings suggest that FGF1 is anti-inflammatory (Fan et al. 2019); however, the mechanism of anti-inflammatory effects of FGF1 is unclear. Also, FGF1 is shown to lower blood glucose levels in diabetic mice but the mechanism of this action is unknown (Suh et al. 2014). We propose that FGF1’s anti-inflammatory and anti-thrombotic actions may be mediated by blocking inflammatory signals through site 2.

Since WT FGF1 is a potent mitogen and, non-mitogenic FGF1 has been sought.

Deletion of N-terminal residues of FGF1 (amino acid residues 21-27), which lacks nuclear translocation signal, has been shown to be non-mitogenic and still has antiinflammatory activity (Imamura et al. 1990). It has been reported that this N-terminal truncated FGF1 induced angiogenesis (Zou et al. 2020). FGF1 R50E has been well characterized as non-mitogenic FGF1. R50E binds to FGFR and heparin, but not to site 1 of integrins. R50E is defective in inducing integrin-FGF1-FGFR1 ternary complex, and defective in inducing sustained ERK1/2 activation and AKT activation (Mori et al. 2008). We showed that non-mitogenic FGF1 R50E (defective in site 1 binding) suppressed tumorigenesis and angiogenesis in vivo (Mori et al. 2013). This indicates that FGF1 R50E has potential as a therapeutic for inflammatory and thrombotic diseases, and insulin resistance.

